# A tetrameric assembly of saposin A: increasing structural diversity in lipid transfer proteins

**DOI:** 10.1101/2021.08.26.457812

**Authors:** Maria Shamin, Samantha J. Spratley, Stephen C. Graham, Janet E. Deane

**Affiliations:** Cambridge Institute for Medical Research, University of Cambridge, Cambridge CB2 0XY, UK; Department of Pathology, University of Cambridge, Cambridge CB2 1QP, UK

**Keywords:** saposin, SapA, lipoprotein, nanodiscs

## Abstract

Saposins are lipid transfer proteins required for the degradation of sphingolipids in the lysosome. These small proteins bind lipids by transitioning from a closed, monomeric state to an open conformation exposing a hydrophobic surface that binds and shields hydrophobic lipid tails from the aqueous environment. Saposins form a range of multimeric assemblies to encompass these bound lipids and present them to hydrolases in the lysosome. This lipid-binding property of human saposin A has been exploited to form lipoprotein nanodiscs suitable for structural studies of membrane proteins. Here we present the crystal structure of a unique tetrameric assembly of murine saposin A produced serendipitously, following modifications of published protocols for making lipoprotein nanodiscs. The structure of this new saposin oligomer highlights the diversity of tertiary arrangement that can be adopted by these important lipid transfer proteins.

## Introduction

Saposins are lysosomal lipid transfer proteins ubiquitously expressed in vertebrates (Hazkani-Covo et al., 2002). The four saposin proteins, named saposin A, -B, -C, and -D (SapA-D), originate from a single precursor protein, prosaposin, that is cleaved within the lysosome. Saposins are required for sphingolipid degradation by lysosomal hydrolases (Kolter and Sandhoff, 2010); in particular, SapA acts as a co-factor for the enzyme β-galactosylceramidase (GALC) by solubilising its galactosphingolipid substrates (Hill et al., 2018; Matsuda et al., 2001; Spiegel et al., 2005). Saposins are small (approximately 80 amino acids, 8-12 kDa) non-enzymatic proteins. Each saposin possesses six conserved cysteines and a conserved N-glycosylation site (Kishimoto et al., 1992). Although individual saposins have low sequence identity (less than 35%), they adopt a common fold consisting of four amphipathic α-helices (Fig. 1A). The N- and C-terminal helices α1 and α4 form a stem stabilised by two disulfide bonds. The central α-helices α2 and α3 form a hairpin motif maintained by a single disulfide bond. Two flexible hinge loops separate the stem and hairpin motifs, allowing saposins to open and close in a jack-knife manner as demonstrated in several structures determined for each saposin (Ahn et al., 2003a; Ahn et al., 2006; Gebai et al., 2018; Hawkins et al., 2005; Hill et al., 2018; Hill et al., 2015; Popovic et al., 2012; Popovic and Prive, 2008; Rossmann et al., 2008). Saposin monomers adopt a closed, globular conformation burying a large hydrophobic surface (Fig. 1B), while the open conformation exposes this surface resulting in dimerisation and encapsulation of lipids and detergents. For SapA, these dimeric assemblies have been captured both in isolation (Fig. 1C) (Popovic et al., 2012) and in the presence of a partner hydrolase (Fig. 1D) (Hill et al., 2018).

**Figure 1.**
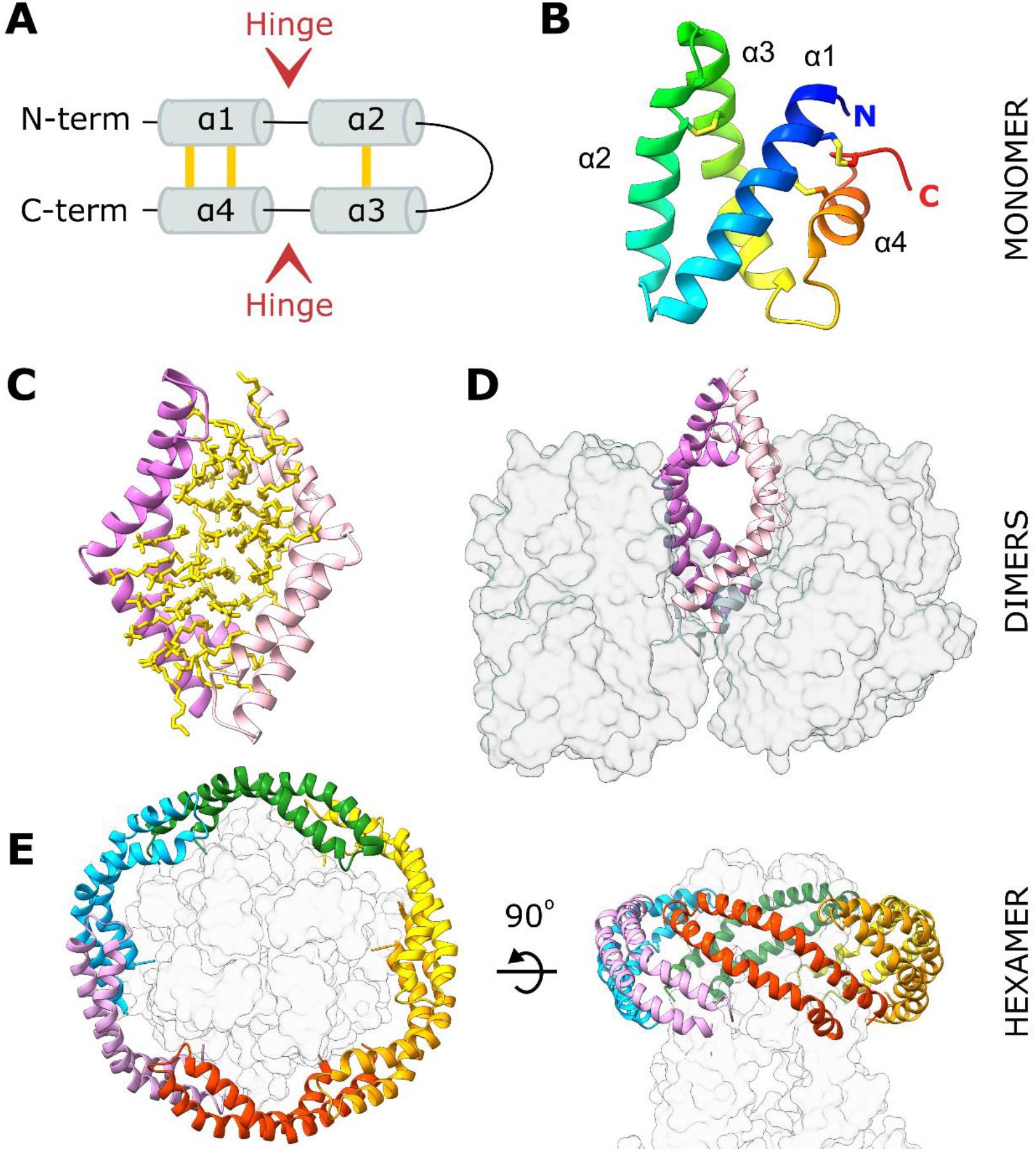
SapA adopts a range of conformations and oligomeric states. **A**. Topology diagram of the saposin fold. Disulfide bonds are shown in yellow. **B**. Crystal structure of a closed human SapA monomer coloured from blue at the N-terminus to red at the C-terminus (PDB ID: 2DOB) (Ahn et al., 2006). **C**. Crystal structure of a human SapA dimer (pink and purple ribbons) encompassing ordered detergent molecules (yellow sticks, PDB ID: 4DDJ) (Popovic et al., 2012). **D**. Crystal structure of a murine SapA dimer (pink and purple ribbons) bound to two molecules of its partner enzyme GALC (transparent grey surface, PDB ID: 5NXB) (Hill et al., 2018). **E**. Cryo-EM structure of a mitochondrial calcium uniporter (transparent grey surface) surrounded by six SapA molecules (coloured ribbons) stabilised by lipids (not illustrated; PDB ID: 6D80) (Nguyen et al., 2018).

The lipid-solubilising properties of SapA have been exploited to form protein-lipid nanodiscs suitable for the reconstitution of transmembrane proteins in a near-native lipid environment for nuclear magnetic resonance (Chien et al., 2017) and electron microscopy (Frauenfeld et al., 2016) studies. These nanodiscs consist of several SapA molecules, in an open conformation, encapsulating lipids surrounding the transmembrane region of these proteins (Fig. 1E) (Flayhan et al., 2018; Nguyen et al., 2018). In this study, we attempted to produce SapA lipoprotein nanodiscs for lipid-binding assays and structural studies with lysosomal hydrolases. These applications required some minor changes to published protocols for nanodisc formation (Chien et al., 2017; Frauenfeld et al., 2016). Interestingly, rather than producing nanodics, X-ray crystallography revealed that SapA assembled into a novel tetrameric arrangement, highlighting the structural diversity of SapA oligomers.

## Materials and Methods

### Saposin A expression and purification

Mouse SapA was expressed and purified as described previously (Hill et al., 2018). Briefly, untagged SapA was expressed in *Escherichia coli* Origami (DE3) cells and bacterial pellets were lysed in anion-exchange buffer (50 mM Tris pH 7.4, 25 mM NaCl). The presence of three disulfide bonds per saposin molecule confers remarkable thermal stability, which has previously been exploited for protein purification without affinity tags (Ahn et al., 2003b; Ahn et al., 2006). Cleared lysate was heat-treated (100 °C, 5 min, repeated four times) and precipitated proteins were cleared by centrifugation. The supernatant containing heat-resistant proteins including SapA was dialysed overnight in the presence of 20 μg/mL DNAse against 5 L of anion-exchange buffer. SapA was further purified by anion exchange chromatography (HiTrap Q Sepharose column) and size-exclusion chromatography (SEC; HiLoad 16/600 Superdex 75 column) equilibrated in 50 mM Tris pH 7.4, 150 mM NaCl. Fractions containing SapA were pooled, concentrated to 16.2 mg/mL and stored at 4 °C. Purified SapA was stable for at least six months.

### Acid Ceramidase (AC) expression and purification

Messenger RNA was extracted from HEK293 cells using an RNeasy Mini kit (Qiagen) according to the manufacturer’s instructions and used as the template to synthesise complementary DNA (cDNA) using QuantiTect Reverse transcription kit (Qiagen) according to the manufacturer’s protocol. PCR product encoding human AC (residues 22-395) was produced from this cDNA using primers (5’-GAAACTAGTCAGCACGCGCCGCCGTGG-3’ and 5’-GTGCTTAAGCCAACCTATACAAGGGTCAGGGC-3’). For production of an inducible, stable cell line, this PCR product was cloned, using SpeI and AflII, into our modified version of the piggyBac target protein plasmid, PB-T-H6, encoding an N-terminal secretion signal and C-terminal hexahistidine tag (Shamin et al., 2019). HEK293F cells were triple-transfected with PB-T-H6 containing AC, PB-RN and PBase using a DNA mass ratio of 5:1:1 and transfected cells were selected with geneticin using the established protocol for the piggyBac expression system (Li et al., 2013). This HEK293F cell line was grown in Freestyle293 medium and protein expression induced by addition of 2µg/mL doxycycline. His6-tagged AC was secreted into the medium and purified by nickel affinity chromatography (Qiagen) with elution in 100 mM citrate pH 4.0.

### Nanodisc formation

The protocol for SapA nanodisc preparation was adapted from Chien *et al*. (Chien et al., 2017) and Frauenfeld *et al*. (Frauenfeld et al., 2016). Egg phosphatidylcholine (PC) was purchased from Avanti Polar Lipids. One mg of PC in chloroform was mixed in a 2 mL glass vial and dried under an argon stream to form a lipid film. The lipid film was resuspended by vigorous vortexing in 104 mM citrate pH 4.0, 156 mM NaCl to make a solution with 5 mg/mL lipids. The lipid solution was incubated in a water bath at 37 °C for one hour, with vigorous vortexing every 15 min. This was followed by 10 cycles of sonication in a sonicating water bath for 30 s and vortexing for 15 s. n-Dodecyl-beta-Maltoside (DDM) from a 5% (w/v) stock was then added for a final buffer composition of 0.2% DDM, 100 mM citrate pH 4.0, 150 mM NaCl, and mixed by pipetting. The solution remained cloudy. SapA protein was diluted to 1.2 mg/mL in nanodisc buffer (100 mM citrate pH 4.0, 150 mM NaCl), 385 µg of mouse SapA was added for each milligram of lipid, and the solution was mixed by pipetting. The solution was incubated at 37 °C for one hour, with mixing every 15 min. Twenty microlitres of nanodisc buffer was then added to make a final volume of 550 µL. The solution was incubated at room temperature for a further 10 min, passed through a 0.2 µm centrifugal filter (Generon) and injected onto a Superdex 200 10/300 column (Cytiva) equilibrated with nanodisc buffer. Fractions containing nanodiscs were concentrated in centrifugal concentrators (4 mL 3K MWCO, then 500 µL 5K MWCO; Amicon and Vivaspin). Nanodiscs were used in experiments within 24 hours. One mg of lipids yielded approximately 400-600 µg of nanodiscs.

### Size exclusion chromatography coupled with multi-angle light scattering (SEC-MALS)

Peak fractions following SEC containing SapA-PC nanodiscs were pooled and concentrated for analysis by SEC-MALS. Sample was injected at room temperature onto a Superdex 200 Increase 10/300 GL column pre-equilibrated in 100 mM citrate pH 4.0, 150 mM NaCl at a flow rate of 0.5 mL/min. The static light scattering, differential refractive index, and UV absorbance at 280 nm were measured in-line by DAWN 8+ (Wyatt Technology), Optilab T-rEX (Wyatt Technology), and Agilent 1260 UV (Agilent Technologies) detectors, respectively. The molar masses were calculated using the protein conjugate analysis algorithm within the ASTRA 6.1 software (Wyatt Technology) and using dn/dc and extinction co-efficients for mSapA calculated using SEDFIT (Schuck, 2000).

### Crystallisation

SapA nanodiscs were used to perform co-crystallisation experiments with the enzyme AC. SapA nanodiscs were concentrated in 100 mM citrate pH 4.0, 150 mM NaCl using a Corning Spin-X UF centrifugal concentrator until SapA was at a concentration of 1.76 mg/mL based on protein absorbance at 280 nm. AC was concentrated to 4.5 mg/mL in 100 mM citrate pH 4.0. Equal volumes of SapA nanodisc and AC solutions were mixed at room temperature. Crystallization experiments were immediately prepared in 96-well nanolitre-scale sitting drops (200 nL protein plus 200 nL of precipitant) equilibrated at 293 K against 80 μL reservoirs of precipitant. A diffraction-quality crystal was grown against reservoir containing 23.25% (w/v) PEG 3350, 0.1 M Bis-Tris pH 5.6. The crystal was cryoprotected with reservoir solution supplemented with 25% (v/v) xylitol and flash-cooled in liquid nitrogen.

### X-ray data collection and structure determination

Diffraction data were recorded on beamline I04 at Diamond Light Source using a Pilatus 6M detector (Dectris). Data were collected at 100 K and λ = 0.9795 Å. Data were indexed and integrated using DIALS (Winter et al., 2018) as implemented by the xia2 processing pipeline (Winter, 2010). Due to significant anisotropic diffraction, diffraction data were subjected to anisotropic scaling using STARANISO (Tickle et al., 2018) and AIMLESS (Evans and Murshudov, 2013). Data collection and processing statistics are detailed in Table 1. The structure was solved via molecular replacement with Phaser-MR (McCoy et al., 2007) by searching for four copies of the open human SapA monomer (Popovic et al., 2012) (PDB ID: 4DDJ). Inspection of electron density maps confirmed the correct placement of three copies of SapA (chains A, B and C) but the position of the fourth SapA chain was clearly incorrect. The fourth molecule was able to be positioned manually exploiting the two fold symmetry of the tetramer, by superposing chain A onto chain C and copying the original chain B to form a new chain D. Following rigid body refinement the tetrameric assembly was confirmed. Refinement was performed iteratively using WinCoot (Emsley et al., 2010), phenix.refine (Afonine et al., 2012) and real-time molecular dynamics-assisted model building and map fitting with the program ISOLDE (Croll, 2018). Refinement was carried out using four-fold non-crystallographic symmetry (NCS) restraints and the ISOLDE model as a reference model, an approach recommended for low resolution structures. The quality of the model was monitored throughout the refinement process using ISOLDE validation tools and Molprobity (Chen et al., 2010). Final refinement statistics are shown in Table 1. The atomic coordinates and structure factors have been deposited in the PDB (Berman et al., 2000) under accession code 7P4T.

**Table 1.**
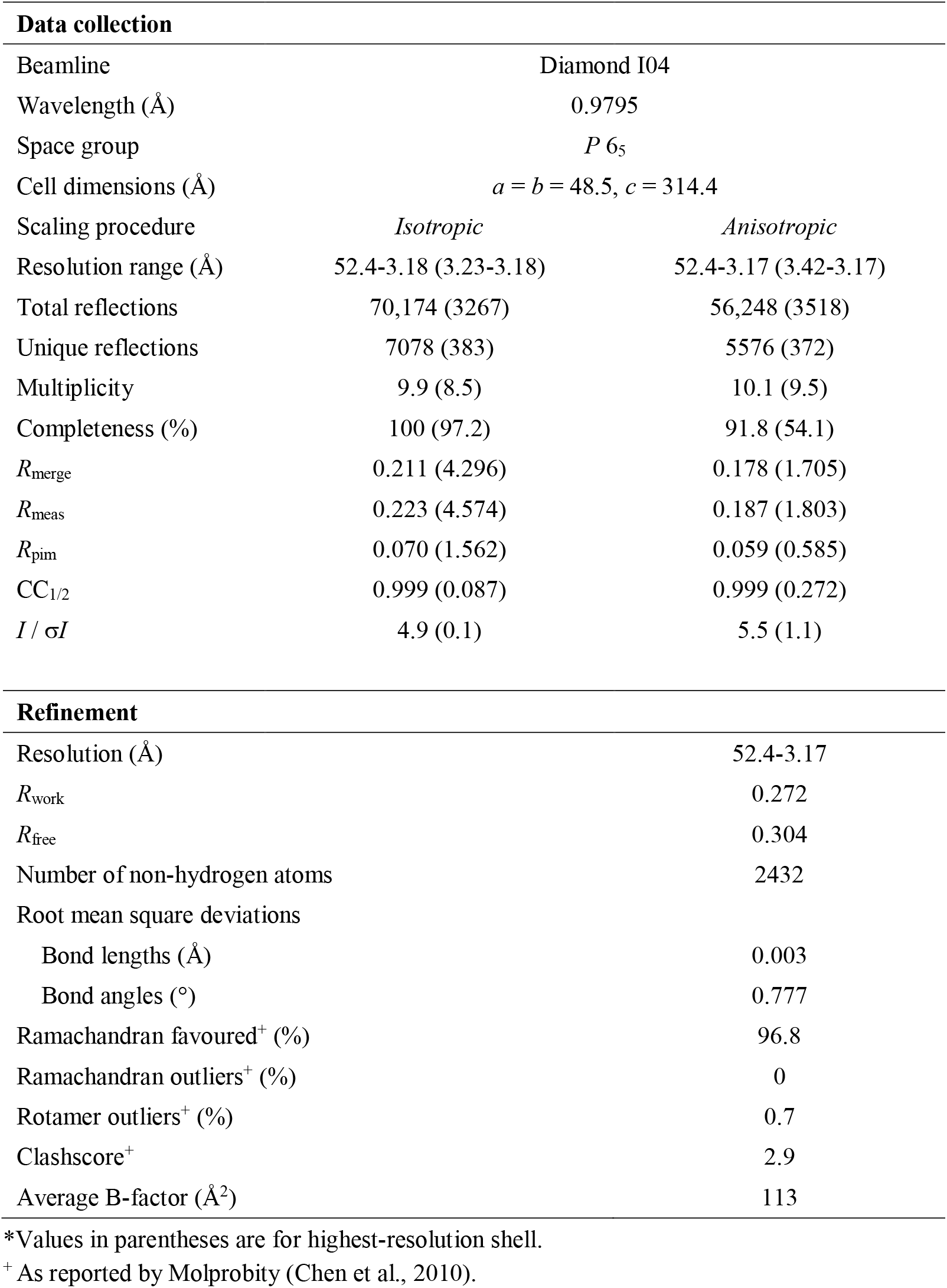
SapA tetramer data collection and refinement statistics.

### Sequence and structure analysis

Multiple sequence alignments were performed using MULTALIN (Corpet, 1988) and visualised with Jalview (Waterhouse et al., 2009). Structure alignments were performed using Secondary Structure Matching or the LSQ algorithm within WinCoot (Emsley et al., 2010). Interface interactions including hydrogen bonds and hydrophobic interactions were identified using PDBePISA (Krissinel and Henrick, 2007). Surface residue hydrophobicity colouring was performed with the Molecular lipophilicity potential tool (Ghose et al., 1998; Laguerre et al., 1997) implemented within ChimeraX (Pettersen et al., 2021). Structural figures were prepared using ChimeraX (Pettersen et al., 2021) and PyMol (Schrödinger, LLC).

## Results

SapA-lipid nanodisc formation requires incubation of SapA and lipids at low pH (pH 4.8) or solubilisation of lipids with the gentle detergent DDM prior to incubation with SapA at neutral pH (Chien et al., 2017). SapA nanodiscs can be formed with a very wide range of lipids (Flayhan et al., 2018) but most studies of this system have employed phosphatidylcholine (PC) (Chien et al., 2017; Flayhan et al., 2018; Leney et al., 2015; Li et al., 2016; Popovic et al., 2012), which was also chosen for this study. The number of SapA and lipid molecules incorporated into the nanodisc and overall nanodisc size depend on several factors, including the lipid to SapA molar ratio (Chien et al., 2017) and the pH of the buffer (Li et al., 2016).

We aimed to use SapA-PC nanodiscs in lipid-binding assays and co-crystallisation experiments with the lipid-processing enzyme acid ceramidase (AC). For these applications, the published protocols (Chien et al., 2017; Frauenfeld et al., 2016) were adapted as follows. Although human SapA has been used to form nanodiscs in previous reports, here we employed the mouse ortholog that our laboratory has used successfully in structural studies of SapA in complex with its partner hydrolase GALC (Hill et al., 2018). Murine SapA possesses 80% sequence identity to human SapA and is essentially identical structurally (RMSD 0.85 Å over 79 C^α^ atoms for the open SapA molecules in PDB 4DDJ and 5NXB). Nanodiscs were prepared in pH 4.0 buffer, close to the low pH of 4.8 used to promote nanodisc formation in Chien *et al*. (Chien et al., 2017). This lower pH was selected as previous work has shown that the interaction of AC with lipid bilayers is strongest at pH 4.0 (Linke et al., 2001). Previous publications reported complete solubilisation of dried lipids in aqueous buffer upon addition of the detergent DDM (Chien et al., 2017; Frauenfeld et al., 2016). However, in our initial experiments lipid solubilisation was incomplete, resulting in a very low final yield. We improved lipid solubilisation by making liposomes in aqueous buffer by 10 cycles of sonication and vortexing of the lipid solution prior to the addition of DDM. Solubilised lipids were then incubated with SapA following the protocol by Frauenfeld *et al*. (Frauenfeld et al., 2016) with a molar ratio of PC to SapA of 30:1 as in Chien *et al*. (Chien et al., 2017). SapA-PC nanodiscs eluted at 15.8 mL from a Superdex 200 10/300 analytical size-exclusion column (Fig. 2A). This elution volume was consistent across several preparations. For initial evaluation of the composition of these nanodiscs, we compared the size exclusion chromatography profile with that from Chien *et al*. (Chien et al., 2017). In this study they demonstrated that using a SapA:lipid ratio of 1:30 resulted in large nanodiscs eluting at 13.3 mL versus a ratio of 1:5 producing smaller discs eluting at 15.9 mL. While we used a lipid:SapA ratio of 30:1, which should have produced a large nanodisc assembly eluting at 13.3 mL, the elution volume of our nanodiscs more closely matched that of a smaller nanodisc assembly produced using a 1:5 ratio and eluting at 15.9 mL. Using size exclusion chromatography coupled to multi-angle light scattering (SEC-MALS), Li *et al*. (Li et al., 2016) showed that nanodiscs eluting at this volume contain two SapA molecules and 23-29 PC molecules. To better characterise the SapA-PC nanodiscs we had produced we also carried out SEC-MALS (Fig. 2B).

**Figure 2.**
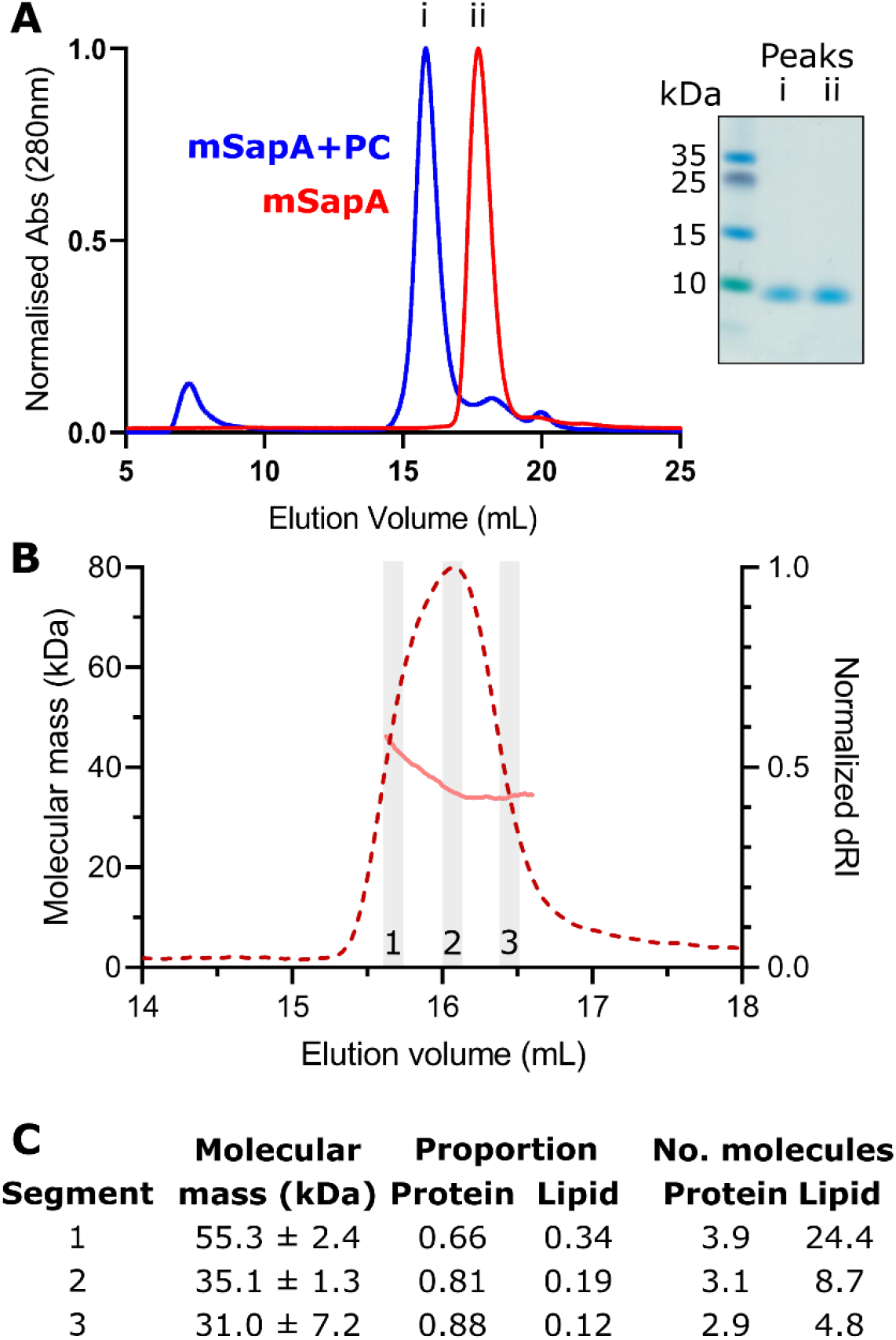
SapA-phosphatidylcholine (PC) nanodiscs preparation. **A**. Overlay of size exclusion chromatography (SEC) elution profiles of (i) SapA following incubation with detergent-solubilised PC and (ii) SapA alone. Peaks eluted at 15.8 mL and 17.7 mL, respectively. Inset: Coomassie-stained SDS-PAGE gel of peak fractions. **B**. Multi-angle light scattering following SEC (SEC-MALS) analysis of SapA-PC assemblies at pH 4.0. The molecular mass distribution (pink line) is shown across the elution profile (normalized differential refractive index, dRI, red dotted line) from a Superdex 200 10/300 Increase column. Data from three regions (grey boxes and numbered 1-3) were used for mass and composition analysis. **C**. Calculated molecular mass and protein:lipid composition across the SEC-MALS peak in (B).

The average molecular mass across the SapA-PC elution peak at pH 4.0 was 36.5 ± 0.4 kDa calculated using a dn/dc value of 0.185 mL/g typical for standard protein samples. However, the sample was clearly heterogeneous, as indicated by higher molecular masses observed at the beginning of the peak and the calculated polydispersity value (Mw/Mn) of 1.03. This heterogeneity may be caused by the presence of a range of nanodisc structures in this sample containing different numbers of SapA and PC. We therefore carried out protein conjugate analysis, whereby the proportion of lipid versus protein can be estimated. For this analysis we used a dn/dc of 0.184 mL/g and UV extinction co-efficient of 0.966 mL/mg.cm for mSapA, calculated using SEDFIT, and a modifier dn/dc of 0.164 mL/g (Inagaki et al., 2013) and UV extinction co-efficient of zero for PC. We carried out this analysis using data from the beginning, centre and end of the observed peak to capture the range of species present in this sample (Fig. 2C). This analysis estimated a range of overall masses from 31 to 55 kDa containing approximately 3-4 molecules of SapA and 5-24 molecules of PC. Although this represents considerable heterogeneity in the nanodisc preparation it is consistent with the mass distribution observed by Li *et al*. (Li et al., 2016) of 35 to 50 kDa, although in that work this was interpreted as a SapA dimer with 23-29 PC molecules.

Having produced SapA nanodiscs possessing a mass consistent with previous studies, we used these in co-crystallisation studies with the lysosomal hydrolase AC with the aim of capturing a snapshot of how this hydrolase functions at the membrane surface. Crystals appeared within 24 hours in 12 different conditions. Three of these conditions produced diffraction-quality crystals, all possessing the same space group and unit cell dimensions. Attempts to solve the structure by molecular replacement with AC failed but a solution was determined using SapA alone. Automated placement of SapA was successful for three molecules of SapA and inspection of maps indicated the presence of a fourth molecule that was placed manually by exploiting the symmetry of the tetramer. Refinement of this tetrameric assembly of SapA revealed that although the fourth molecule was not as well ordered as the other three, there was clear evidence of its orientation and conformation in the electron density maps. Despite AC being present in the crystallisation drops there was no evidence of this protein in the crystal structure.

The tetrameric structure of mouse SapA captured here is a distinct assembly from previously described SapA nanodiscs or other saposin oligomers. Four SapA molecules in an open conformation form a diamond-shaped assembly possessing three orthogonal two-fold rotation axes (Fig. 3A). This symmetry is not perfect as the individual monomers possess small differences in the conformations of loop regions and the C-terminal helix (Fig. 3B, RMSDs 0.8 – 1.4 Å^2^ over 80 C^α^ residues), resulting in non-crystallographic symmetry of the tetramer. This lack of symmetry is best illustrated by the interface between chains A and C where N77 engages in a hydrogen bond with the backbone of S74, whereas chains B and D do not form this interaction (Fig. 3A, right panel and Fig. 3C). This SapA tetramer is maintained via several protein-protein contacts between SapA monomers including both hydrophobic and polar interactions. Each chain of the tetramer interacts directly with the three other chains burying a total surface area of 1210 Å^2^ for each monomer, representing approximately 20% of the SapA surface (Fig. 3D). The residues participating in these interactions are distributed throughout the entire SapA sequence and are mostly conserved between mouse and human SapA (Fig. 3E) suggesting that this novel tetrameric assembly is not driven by interactions specific to mouse SapA.

**Figure 3.**
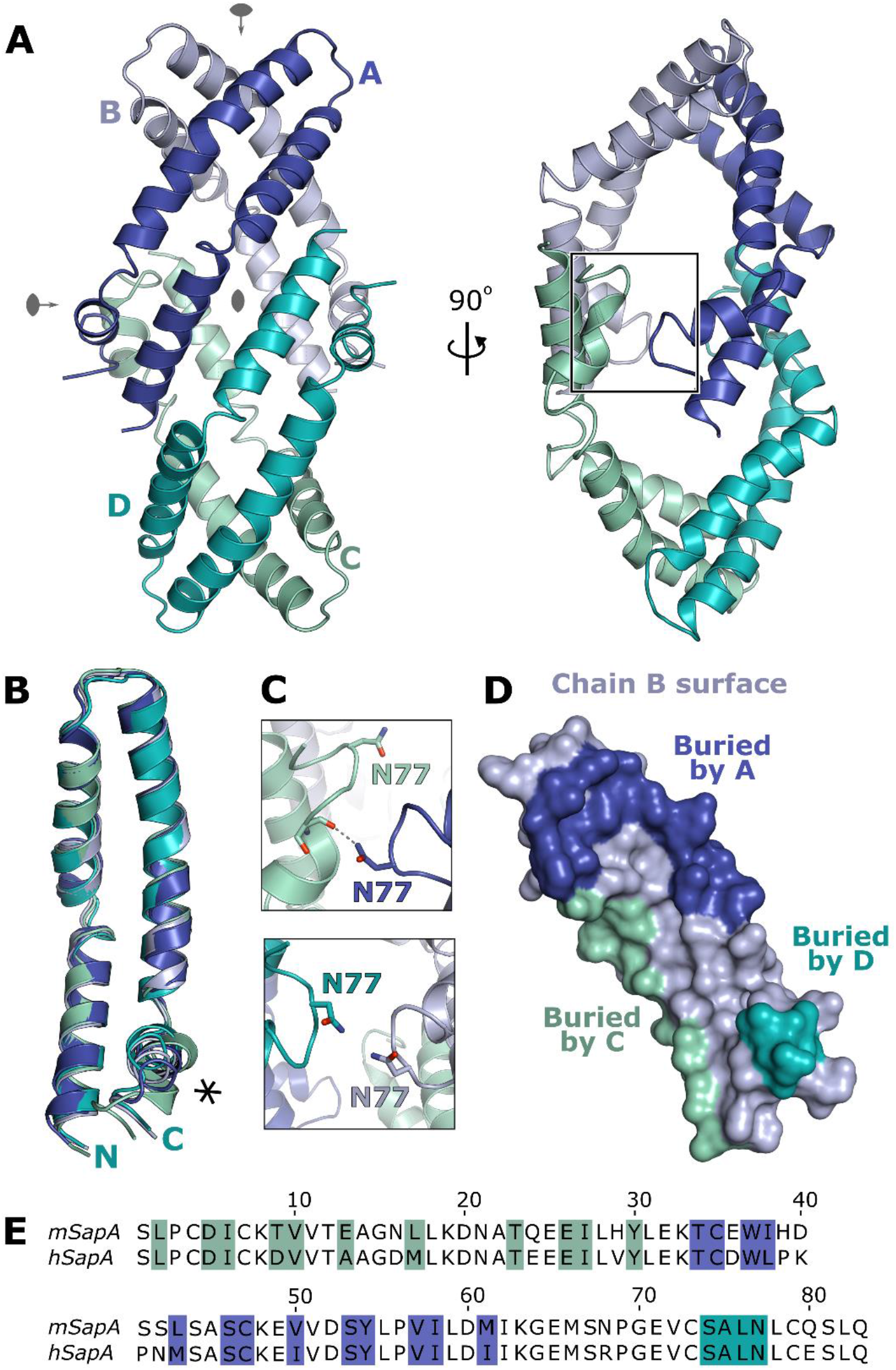
Crystal structure of the tetrameric assembly of SapA. **A**. Ribbon diagram of the SapA tetramer (chains A-D labelled and coloured blue, light blue, cyan and green cyan) illustrating the three orthogonal quasi-two-fold rotation axes (grey symbols). Two views of the tetramer are shown rotated by 90° to illustrate the overall assembly. The region detailed in panel C is boxed. **B**. Overlay of the four SapA chains in the assembly illustrating the structural changes near the C-terminus (asterisk). **C**. Comparison of the equivalent interfaces between chains A and C (top panel) and chains B and D (bottom panel) illustrating a break in the symmetry of the tetramer. **D**. Surface representation of the B chain of the tetramer, in the same orientation as in panel A, with residues involved in intermolecular interactions coloured according to the interacting chain (colouring as in A). **E**. Sequence alignment of mouse SapA (mSapA) and human SapA (hSapA) showing that the contact residues are highly conserved (coloured as in panel D).

Although the tertiary arrangement of SapA determined here is substantially different from previously observed assemblies of SapA (Fig. 1), the open conformation of the SapA monomers within these structures are very similar (Fig. 4A). Human SapA within the nanodisc structure has a RMSD from mouse SapA within the tetramer of 1.1 Å (over 80 C^α^ atoms) and GALC-associated mouse SapA has an RMSD of 1.5 Å (over 80 C^α^ atoms). This similarity allows direct comparison of the different tertiary assemblies. The nanodisc assembly (Fig. 1C) (Popovic et al., 2012) possesses two human SapA molecules enclosing an ordered bilayer of the detergent lauryldimethylamine oxide (LDAO). In this structure, the entire assembly is held together by interactions between lipid molecules and the hydrophobic acyl core formed by two SapA molecules arranged in a head-to-tail conformation with no direct contacts between SapA monomers. The equivalent dimer within the tetramer structure is very different, possessing a head-to-head arrangement of monomers and substantial direct protein-protein interactions (Fig. 4B). Interestingly, this head-to-head arrangement is more similar to the structure of mouse SapA in complex with the lipid processing enzyme GALC (Fig. 4C) (Hill et al., 2018). Both structures are maintained by intermolecular hydrophobic contacts near the loop formed by residues 38-42; however, the residues participating in these contacts and the position of the SapA monomers relative to each other differs. These differences result in intermolecular clashes when the tetramer is docked onto the GALC surface and there is no opening over the GALC active site to allow lipid processing. Comparison of these structures suggests that the tetrameric assembly of SapA is not compatible with binding to GALC without some structural rearrangement.

**Figure 4.**
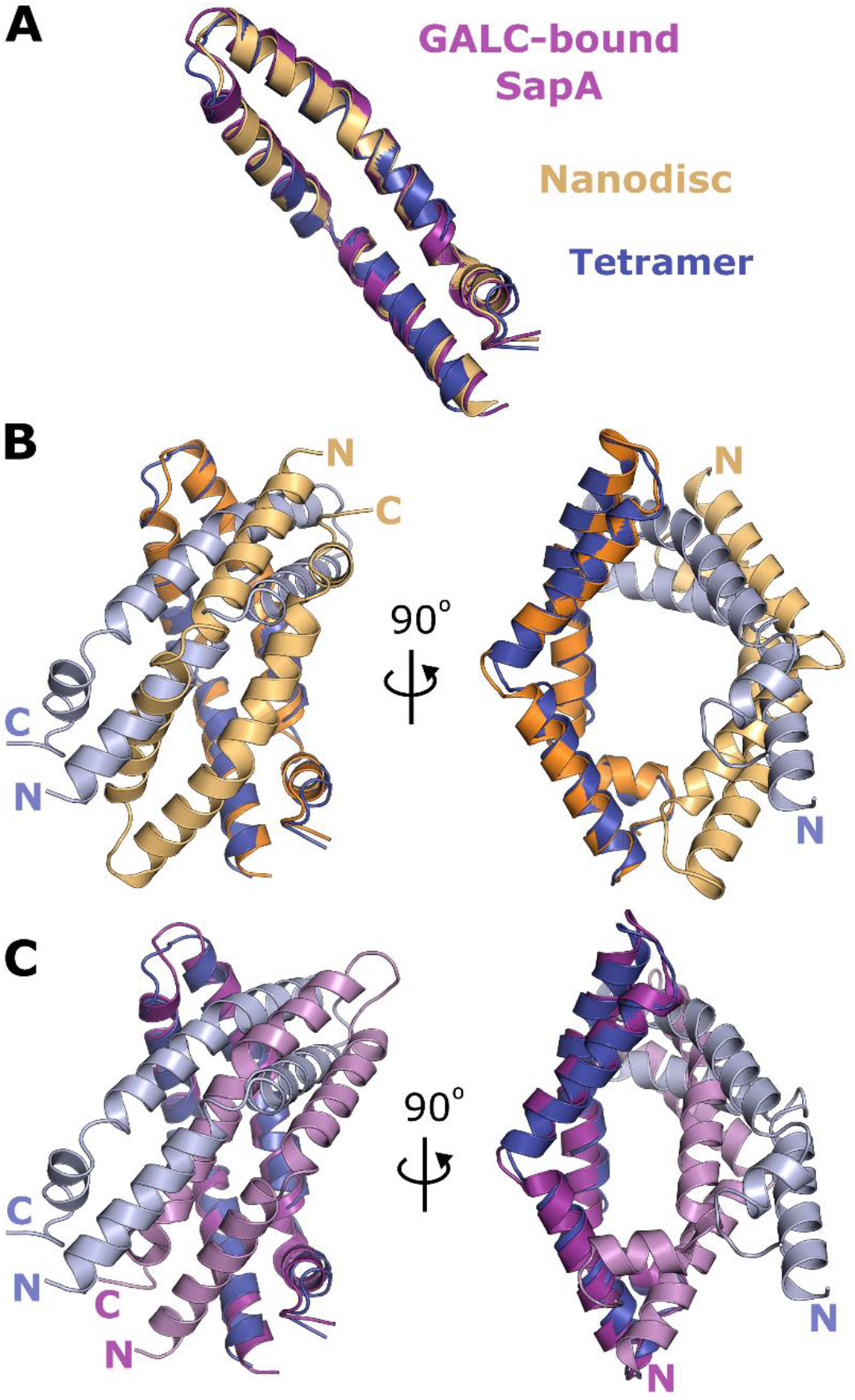
Comparison of SapA oligomers. **A**. Overlay of the open conformation of SapA monomers in the tetrameric structure (blue), GALC-associated dimeric SapA (purple) (Hill et al., 2018) and the nanodisc dimer (orange) (Popovic et al., 2012). **B**. Overlay of the SapA dimer in the nanodisc structure (orange and light orange) with two chains of the SapA tetramer (blue and light blue). **C**. Overlay of the SapA dimer present in the GALC complex (purple and pink) with two chains of the SapA tetramer (blue and light blue). Two views rotated by 90° are shown for both panels B and C.

The core of the SapA tetramer structure is hollow and potentially solvent-accessible if it is not filled with amphipathic molecules (Fig. 5A). Considering the highly hydrophobic nature of the internal SapA surface (Fig. 5B), exposure of this hollow core to solvent would be thermodynamically unfavourable. It is therefore likely that disordered PC and/or DDM molecules are present in this internal cavity. Indeed, there is observable electron density in the cavity suggesting the presence of lipid or detergent head groups in this space. However the weakness of this density suggests these molecules are not well ordered and they were therefore unable to be modelled accurately in this structure.

**Figure 5.**
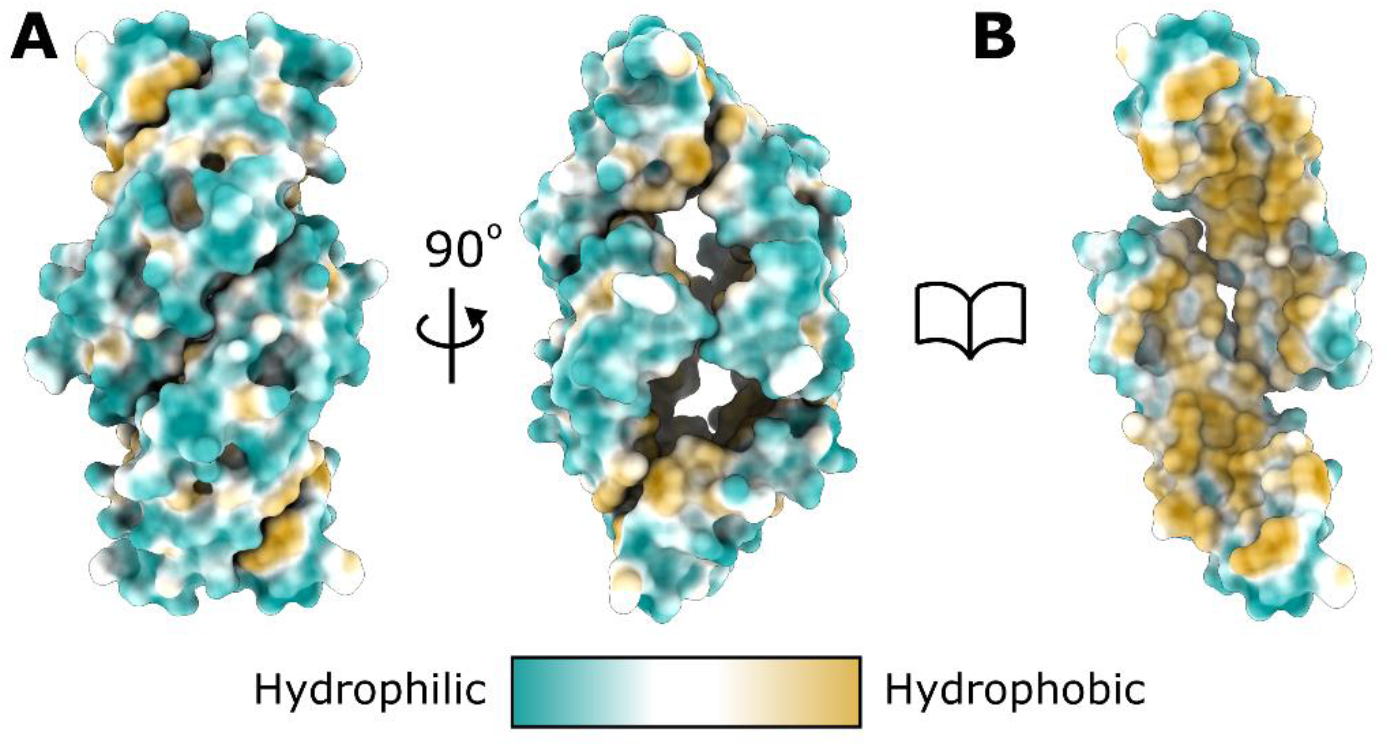
The core of the SapA tetramer is highly hydrophobic. **A**. Surface representation of the SapA tetramer coloured by residue hydrophobicity. The structure is displayed in the same orientations as shown in Fig. 3A. **B**. The tetramer with two SapA chains removed, oriented as in the left panel, reveals the core of the tetramer to be highly hydrophobic.

## Discussion

In this study, we show that SapA incubated with detergent and lipids can form a tetrameric assembly differing from previously observed SapA lipoprotein nanodiscs (Chien et al., 2017; Popovic et al., 2012). The production of a tetrameric SapA assembly instead of SapA nanodiscs could be explained by two differences between our protocol and the original protocols for nanodisc formation. 1) Previous protocols use human SapA for nanodisc formation, whereas mouse SapA was used in this study. However, these orthologs are highly similar, and the residues involved in the interactions maintaining this tetrameric assembly are identical or similar in their hydrophobic properties between human and mouse SapA. It therefore seems unlikely that this modification accounts for the difference in the final product. 2) In order to have the best chance of capturing a complex of AC with a lipid bilayer, we prepared SapA nanodiscs at pH 4.0, whereas previous studies were performed at pH 4.8 to 7.5 (Chien et al., 2017; Frauenfeld et al., 2016; Li et al., 2016). Li *et al*. (Li et al., 2016) observed that the oligomeric state of SapA within nanodiscs is dependent on the final pH of the nanodisc solution, rather than the pH at which these were formed. This supports the idea that SapA assemblies are highly dynamic, adopting distinct oligomeric states depending on the buffer conditions.

The SEC-MALS analysis of the SapA-nanodisc preparation identified a range of species present in solution including masses consistent with the tetrameric assembly observed in the X-ray crystal structure (Fig. 2C). However, this solution sample also contained masses consistent with a trimeric SapA assembly and are in rough agreement with the masses observed by Li *et al*. (Li et al., 2016) that were interpreted as dimeric SapA nanodiscs containing 23-29 PC molecules. It remains unclear if the heterogenous SapA nanodisc sample underwent structural rearrangements during the crystallisation experiment such that the majority of SapA molecules shifted to the tetrameric form, or whether the tetrameric assembly preferentially formed crystals from within the heterogenous mix of tetramers and nanodiscs. Alternatively, the presence of AC in the mixture may have influenced the conformation of the SapA nanodisc assemblies by interacting with the lipids directly (Linke et al., 2001).

The structure determined here represents a distinct tertiary arrangement of SapA and an oligomeric assembly unseen previously for any of the saposin family members. The ability of SapA to form variable oligomeric assemblies may have relevance for its function as a lipid transfer protein in the lysosome.

## Acknowledgements

We thank Randy Read for assistance with the molecular replacement solution of SapA, in particular helping us place the fourth chain. We acknowledge Diamond Light Source for time on beamline I04 under proposal MX15916. Remote access was supported in part by the EU FP7 infrastructure grant BIOSTRUCT-X (Contract No. 283570). M.S. was supported by a Wellcome Trust PhD studentship [203984]. This work was supported by MRC grant MR/N020626/1 to J.E.D. S.C.G. was supported by a Sir Henry Dale Fellowship co-funded by the Royal Society and Wellcome Trust (098406/Z/12/Z). J.E.D. was supported by a Royal Society University Research Fellowship (UF100371) and a Wellcome Trust Senior Research Fellowship (219447/Z/19/Z).

## Notes

### Competing Interest Statement

The authors have declared no competing interest.

